# Hierarchical whole-brain modeling of critical synchronization dynamics in human brain

**DOI:** 10.1101/2024.05.08.593146

**Authors:** Vladislav Myrov, Alina Suleimanova, Samanta Knapič, Paula Partanen, Maria Vesterinen, Wenya Liu, Satu Palva, J. Matias Palva

## Abstract

The brain operates at the critical transition between order and disorder which supports optimal information processing. Whole-brain computational modeling is a powerful tool for uncovering the system-level mechanisms behind large-scale brain activity in both healthy and pathological states. However, most previous approaches have focused on either functional connectivity or criticality, making it difficult to capture both aspects simultaneously. Here, we introduce a new method based on a Hierarchical Kuramoto model that incorporates two levels of hierarchy. In our model, each node contains a large number of coupled oscillators, which allows us to examine both local synchronization and long-distance interactions between brain regions. The model produces critical-like dynamics marked by emergent long-range temporal correlations (LRTCs) and both inter-areal phase synchronization and amplitude correlations during the transition from asynchronous to synchronous states. Notably, structure-function coupling shows distinct patterns: correlations with structural connectivity peak at criticality for LRTCs and amplitude correlations, but decay for local and inter-areal phase synchronization. Comparisons with human resting-state magnetoencephalography (MEG) data reveal that the model’s behavior most closely resembles MEG phase synchronization and multi-peak power spectra on the subcritical side of an extended critical regime, supporting the hypothesis that the human brain operates in this state.

**Significance Statement:** While criticality has gained attention in neuroscience, it is often considered distinct from other emergent properties such as functional connectivity (FC). However, recent experimental evidence suggests that a system’s position within a critical state space governs its dynamics including FC. Here, we introduce a hierarchical modeling framework for whole-brain synchronization dynamics based on local and network-level control parameters. We investigated how the operating point shapes structure-function coupling and spectral properties, and show that model observables best match magnetoencephalography (MEG) data in a near-critical regime, suggesting that the human brain operates in this state. Our work provides a framework for modeling whole-brain-scale activity and supports the view that criticality and classical emergent properties are unified aspects of oscillatory dynamics.

---

**E**lectrophysiological activity is characterized by neuronal oscillations that are rhythmic excitability fluctuations arising with frequency-specific synaptic mechanisms in neuronal micro- and macrocircuits (1). Human brains *in vivo* exhibit moderate levels of long-range phase synchronization of neuronal oscillations in both intra-cerebral stereo-EEG (SEEG) (2–4) and non-invasive magneto- and electroencephalographic (M/EEG) recordings (5–7). Synchronization plays a mechanistic role in regulating neuronal communication in distributed brain networks (8–10) underlying cognitive functions (11–13). In contrast, both hypo- and hypersynchronization constitute core pathophysiological mechanisms in neurological disorders, such as epilepsy (3, 14) and Parkinson’s disease (15), and characterize neuropsychiatric diseases, such as depression (16, 17), where abnormalities in synchrony are correlated with the severity of depressive symptoms (18–20).

Electrophysiological methods such as M/EEG and SEEG in humans (Figure 1A) provide the millisecond-scale temporal resolution needed to observe these oscillations in frequency ranges from 1 to *>*200 Hz. Typically, data from these recordings are filtered to obtain narrow-band analytical time series (Figure 1B) and observables, such as local power spectra (Figure 1C), and amplitude time series are then used as proxy measures for local (within-parcel) neuronal synchronization. Pairwise interareal amplitude correlation (Figure 1D,F) and phase synchronization (Figure 1E,G), or their multi-variate counterparts can be used to assess the functional connectivity (FC) of inter-areal interactions. Neuronal oscillations exhibit remarkable variability across individuals in their power and levels of inter-areal coupling (21). This variability is functionally significant as it explains, *e*.*g*., difference in cognitive performance (16, 22), learning (23), and neurological state (14).

**Fig. 1.**
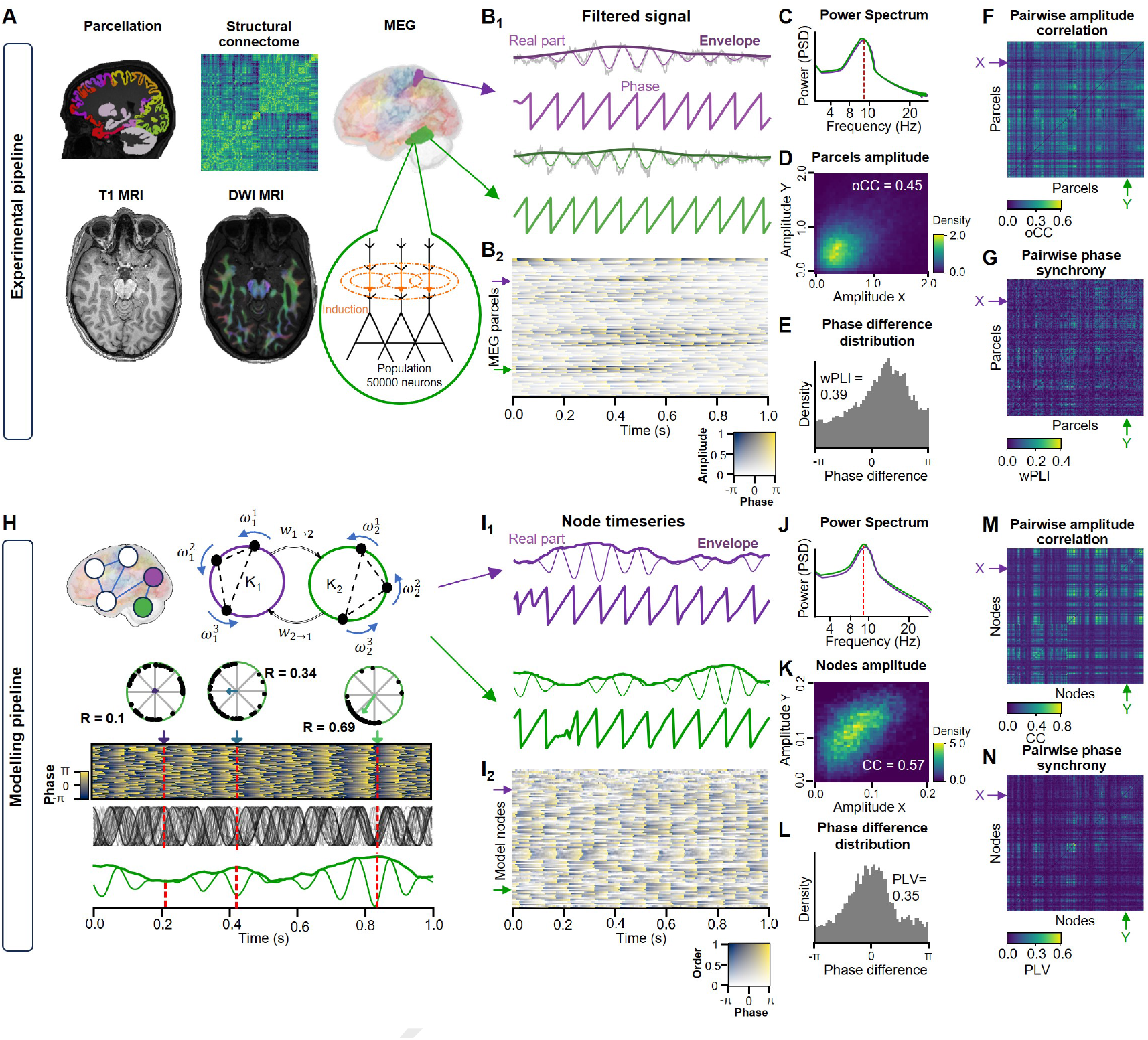
General representation of an experimental pipeline. **A** Summary of imaging and data processing methods, with T1-weighted magnetic resonance imaging (MRI) for anatomical parcellation, Diffusion-weighted imaging (DWI) MRI for structural connectome reconstruction, and MEG for functional data acquisition. *B*_1_ MEG signals from the two representative parcels in (A) showing the broadband signal (gray) overlaid with the narrow-band filtered signal at 10 Hz, its phase and envelope. *B*_2_ Phase heatmap of MEG signals across channels, with low-amplitude regions shown as transparent. **C** Power spectral density (PSD) for the two parcels. **D** Two-dimensional histogram of amplitudes for the 10 Hz parcel signals in (B). **E** Distribution of phase differences between a pair of MEG signals, with a weighted phase lag index (wPLI) of 0.39. **F** Heatmap of pairwise orthogonalized amplitude correlations (oCC) of alpha-band MEG signals (10 Hz). **G** Pairwise wPLI matrix of alpha-band MEG signals (10 Hz). **modeling pipeline. H** On top level, the model is made of multiple inter-connected nodes where each node represents a single brain region. Each node comprises a large amount of Kuramoto oscillators with central frequency *ω* and intra-nodal coupling strength of *K*_*n*_. The complex average across oscillators yields the node timeseries of which the absolute value is node order (R) that is comparable with oscillation amplitude in MEG (see B). *I*_1_ Node time series and corresponding phase *I*_2_ for two model nodes. **J** Power spectral density of the real part of simulated time series. **K** Two-dimensional histogram of amplitudes for two simulated signals. **L** Phase difference distribution for two simulated signals, with an example PLV value of 0.35. **M** Pairwise amplitude correlation heatmap for simulated time series. **N** Pairwise PLV matrix of simulated signals.

The framework of criticality provides a statistical physics approach to understanding these neuronal dynamics and their variability. The “critical brain” hypothesis (24–27) proposes that the operating point (OP) of neuronal systems *in vivo* lies at the phase transition between the subcritical (disordered) and supercritical (ordered) phases in the system’s state space. Operation at this critical point leads to the emergence of scale-free spatiotemporal correlations, observable both as longrange temporal correlations (LRTCs, (26)) and power-law scaled avalanches (27), and yields several functional advantages, such as maximized information capacity (24), complexity (28) and processing, dynamic range (29), and transmission rate (30, 31).

Operating point indicates the system’s position in the state space and is regulated by underlying control parameters that for neuronal systems are primarily thought to be excitation-inhibition (E/I) balance maintained both by fast neurotransmission (32–35) and slow neuromodulation (36, 37), where the ascending reticular arousal system is a potent dynamic E/I modulator (38) as well as structural connectivity (SC) strength between brain areas (39, 40). A given OP is associated with specific emergent dynamics, which can be operationalized with a range of synchronization and criticality observables (21, 26, 27, 33, 41). Because neither the control parameters nor the system’s states are observable from measurement data, their relationships with observables have remained a topic of interest, whereas computational modeling is a useful method for studying the causal effect of system control parameters on its dynamics (42).

Numerous computational models have been developed to investigate the principles underlying whole-brain neural dynamics (43–46). These models are made “whole-brain” relevant by connecting the model’s nodes into a network with weights obtained from the structural connectome of human white-matter connections measured with diffusion tensor spectrum (47). Inter-areal phase synchronization has been modeled, *e*.*g*., with the Wilson-Cowan model (4), neural mass models (48), and the Kuramoto model (49, 50). Likewise, inter-areal amplitude correlations have been addressed with neural mass models (51) and Hopf models (52), and have also been used as a neuronal correlation proxy for functional magnetic resonance imaging (fMRI) functional-connectivity observables (53, 54). In contrast, computational models in criticality research have often focused on avalanches using the branching-process (55) and Ising models (56) from statistical physics, which do not express oscillations. Neurophysiology-inspired population models (57–59) and networks of excitatory and inhibitory neurons, such as the critical oscillations (CROS) model (33), have shown that partially synchronized oscillations may exhibits LRTCs peaking at criticality (60, 61). However, those models represent small neuronal populations rather than capturing whole-brain scale dynamics. There is thus a shortage of whole-brain scale models that capture oscillation-based phase- and amplitude correlations concurrently with critical dynamics.

The Kuramoto model is one of the basic models of synchronization dynamics and captures the emergence of selforganized synchrony in a system of coupled oscillators (62). The model exhibits a phase transition between subcritical (asynchronous) and supercritical (synchronous) phases, which is characterized by emergence of critical-like dynamics, such as LRTCs (21) and avalanches (63). Prior research using Kuramoto modeling in the neuroscience context has represented the activity of neuronal populations within cortical areas with single Kuramoto oscillators coupled by realistic interareal SC (64). This approach yields whole-brain network dynamics but does not offer node-level observables, such as order fluctuations or LRTCs, which are essential for critical dynamics. Another approach for using the Kuramoto model represents a neuronal system as either a single (65, 66) or two large populations (50, 67) of oscillators, which captures critical-like dynamics at a “local” level similarly to the CROS model, but does not extend the model into structural-connectivity defined whole-brain scale networks.

We present here a new hierarchical extension of the classic Kuramoto model. Hierarchical Kuramoto consists of a wholebrain network where each node itself is a system of a large number of oscillators. This multi-level architecture yields separable local and inter-areal synchronization dynamics and allows for direct comparability with local and inter-areal observables in neuroimaging data. We first used Hierarchical Kuramoto to investigate the dependence of structure-function coupling on the operating points. Then, to understand the likely range of operating points in human brain dynamics, we asked where in the parameter space the emergent model dynamics would best match the experimental multi-scale dynamics and power spectra observed with MEG.

## Results

### Whole-brain model of critical oscillations

Brain activity emerges through interactions on both micro- and macroscopic scales from small neuronal assemblies to large neuronal systems observable even with on-scalp electrophysiological methods such as MEG (Figure 1A). Complex interactions across micro to large scale dynamics give rise to oscillatory phenomena with rich phase and amplitude envelope dynamics at different time-scales (Figure 1B), which are measurable, *e*.*g*., with observables such as power spectra (Figure 1C), amplitude correlation (Figure 1D), and phase synchronization (Figure 1E).

To capture intra- and inter-areal synchronization dynamics concurrently and in a dissectable manner, we designed a hierarchical model where each node was a full system of Kuramoto oscillators (1, 50, 62) analogous to each cortical area containing a large population of neurons. The oscillators were directionally coupled within each node with coupling weighted by the local control parameter K (Figure 1H, see Methods). We quantified within-node synchronization, *i*.*e*., ‘node order’ with the Kuramoto order. These local synchronization dynamics, internal to each node, were (Figure 1H and I) similar to the *in vivo* local neocortical synchronization dynamics observable in MEG oscillation amplitude fluctuations (Figure 1B) both phenomenologically and in terms of their power spectra (Figure 1C and J).

On the whole-brain level of hierarchy, we defined the interaction between the nodes with the directional phase-difference of respective complex node time series weighted by node order and strength of white-matter structural connections between the nodes (see Methods). The local synchronization fluctuations (Figure 1I) were paralleled by inter-areal FC in the forms of amplitude (order) correlations (Figure 1K,M) and phase synchrony (Figure 1L,N), again with phenomenological similarity with those observed in MEG data.

### Local and global control parameters shape the model critical-like dynamics

The critical brain hypothesis posits that neural networks *in vivo* operate near a critical point between order and disorder characterized by moderate levels of synhcronization to optimize computational capabilities and adaptability (24, 25). The critical region in the state space emerges from a position in parameter space. In oscillatory network models, these parameters are local and global coupling strengths (Figure 2A). A phase transition occurs between transformation from desynchronization to complete synchronization on both nodal and inter-node levels, which is operationalized with node order and inter-node phase synchronization, respectively. The critical dynamics emerge at the phase transitions where oscillations are characterized by LRTCs, operationalized using Detrended Fluctuation Analysis (DFA, Figure 2B) these peaking at the critical phase transition.

**Fig. 2.**
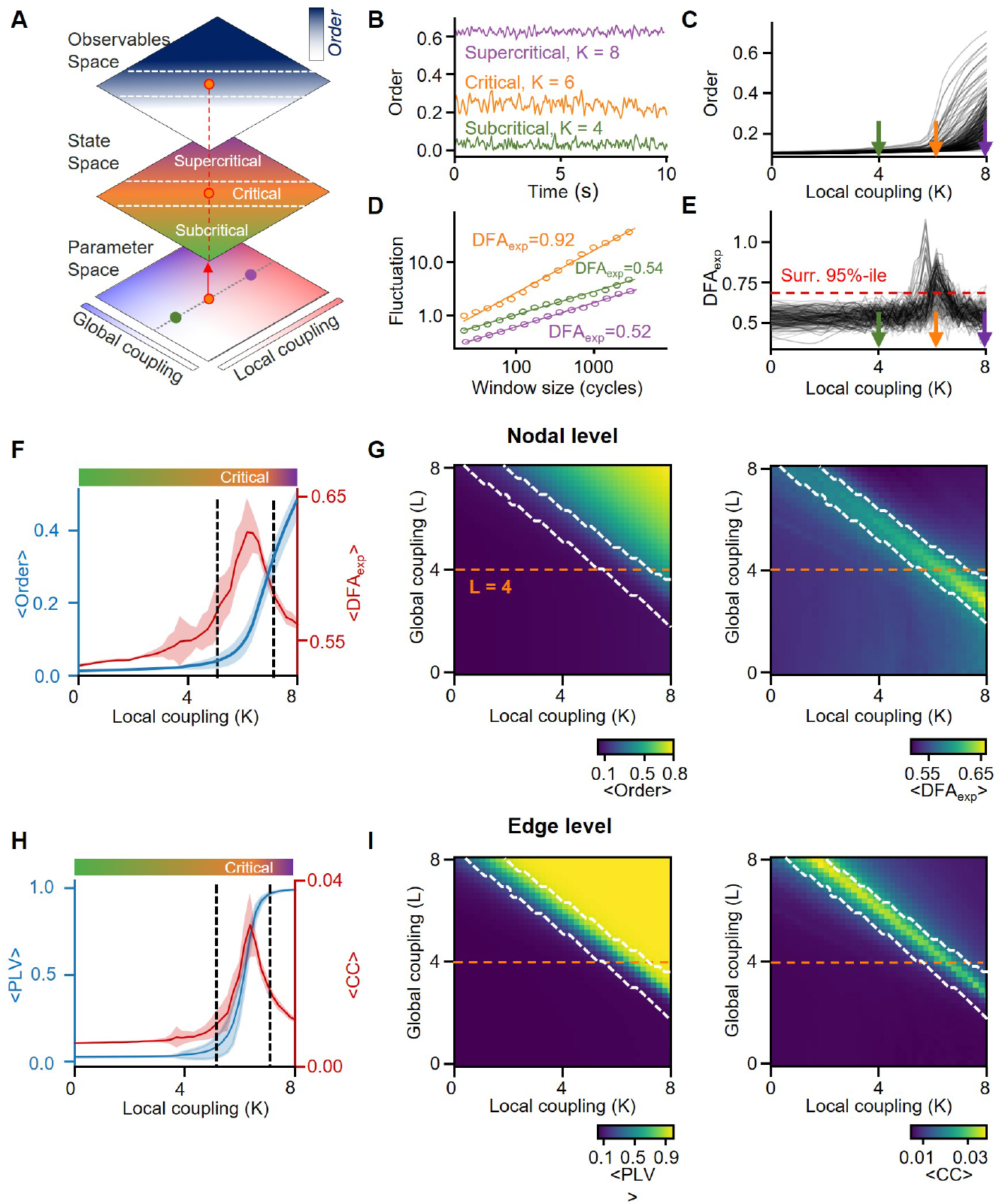
Phase transition and emergence of critical-like dynamics in Hierachical Kuramoto Model. **A** Simplified representation of the parameter space of Hierarchical Kuramoto, illustrating two key control parameters: local coupling strength (K) and global coupling strength (L). Colored dots indicate three distinct parameter combinations giving rise to unique operating points in the subcritical, critical, and supercritical parts of the state space, leading to observables such as synchronization and long-range temporal correlations (LRTCs). **B** Examplar timeseries generated with Hierarchical Kuramoto model from subcritical, critical and supercritical regimes states. **C** An average model order as a function of local coupling strength (K). One line represents a single model node. **D** Detrended Fluctuation Analysis (DFA) fits and their exponents (*DFA*_*exp*_) for each regime. **E** DFA exponent (*DFA*_*exp*_) as a function of local coupling strength (K), where each line represents a single model node. The red dashed line indicates a 95-th percentile of DFA exponent obtained from surrogate data (white noise). **F** Average model order (blue) and *DFA*_*exp*_ (red) as a function of local coupling strength (K). The critical regime (shaded) is defined as the region where more than 10% of nodes exhibit *DFA*_*exp*_ *>* 0.65. Shaded areas represent standard error (SE) confidence intervals (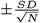, where *N* = 24 models). **G** Heatmaps of average order (left) and *DFA*_*exp*_ (right) across model nodes as functions of local (K) and global (L) coupling strengths. The white contour outlines the critical regime, defined as regions where more than 10% of nodes have *DFA*_*exp*_ *>* 0.65. **H** Average Phase Locking Value (PLV, blue) and cross-correlation (CC, red) as functions of local coupling strength (K). **I** Heatmaps of PLV (left) and CC (right) averaged across edges, as functions of local (K) and global (L) coupling strengths.

To assess the emergence of critical dynamics in the Hierarchical Kuramoto model, we first used a single-subject SC matrix to define inter-areal coupling weights. We then simulated resting-state activity (see Methods) and computed the DFA exponent for each node of the simulated time series as a function of the local coupling coefficient (K).

We found that the hierarchical approach effectively modeled the critical transition at individual node level from low to high local synchronization (Figure 2B,C). This transition featured a peak in the DFA scaling exponent near 1, indicating critical-like dynamics (Figure 2C,D).

Next, we extended the analysis by varying global coupling (L) and local coupling (K) strength simultaneously. We created a cohort of structural-connectome-informed models (*N* = 24) and analyzed the behavior of intra- and inter-node level observables. At the nodal level, we found that the DFA exponent peaks during the phase transition of the node order (Figure 2F). On the KL surface – the DFA exponent as a function of both control parameters – the critical region was represented by a linear ridge and reached a maximum when local coupling was slightly lower than global (Figure 2G). Notably, the inter-subject variance began to grow before the DFA exponent peaked and achieved maximum at at criticality.

Inter-nodal interaction dynamics showed a transition from near zero synchronization to almost perfect synchrony, albeit the cross-correlation between node order timeseries peaking during the transition period and its peak matching the critical region assessed with LRTCS (Figure 2H,I). Unlike the DFA exponent that was maximized when *K > L*, the cross-correlation achieved the highest values when *L > K*.

Similar results were reproduced with a log-transformed structural connectome matrix (see Supplementary Figure 1). However, the variability between critical peaks was lower and the critical ridge was narrower in contrast to the original nontransformed SC. Consequently, the average DFA exponent and CC were higher when using log-transformed edge weights. This highlights that the emergence of LRTCs is not a property of a specific structural architecture but a general phenomenon.

### Interaction-specific breakdown or maximization of structure– function coupling at criticality

The SC is a backbone of functional relations in the brain and previous studies have shown that models initialized with the human structural connectome can reconstruct patterns of resting state networks (40, 68). Although their primary focus was specific features of activity, such as FC, they did not investigate behavior of multiple observables at once.

In this study, we expended the scope beyond phase internodal synchronization. To analyze relationships between the model’s structural architecture and oscillatory dynamics, we computed the Pearson correlation coefficient between inter-node connection weights and observables of oscillatory dynamics. We correlated node strength (average SC of a node) with node-level observables (e.g., order, DFA exponent), and inter-node statistics (e.g., edge strength of PLV/CC) were correlated with edge weights (Figure 3A).

**Fig. 3.**
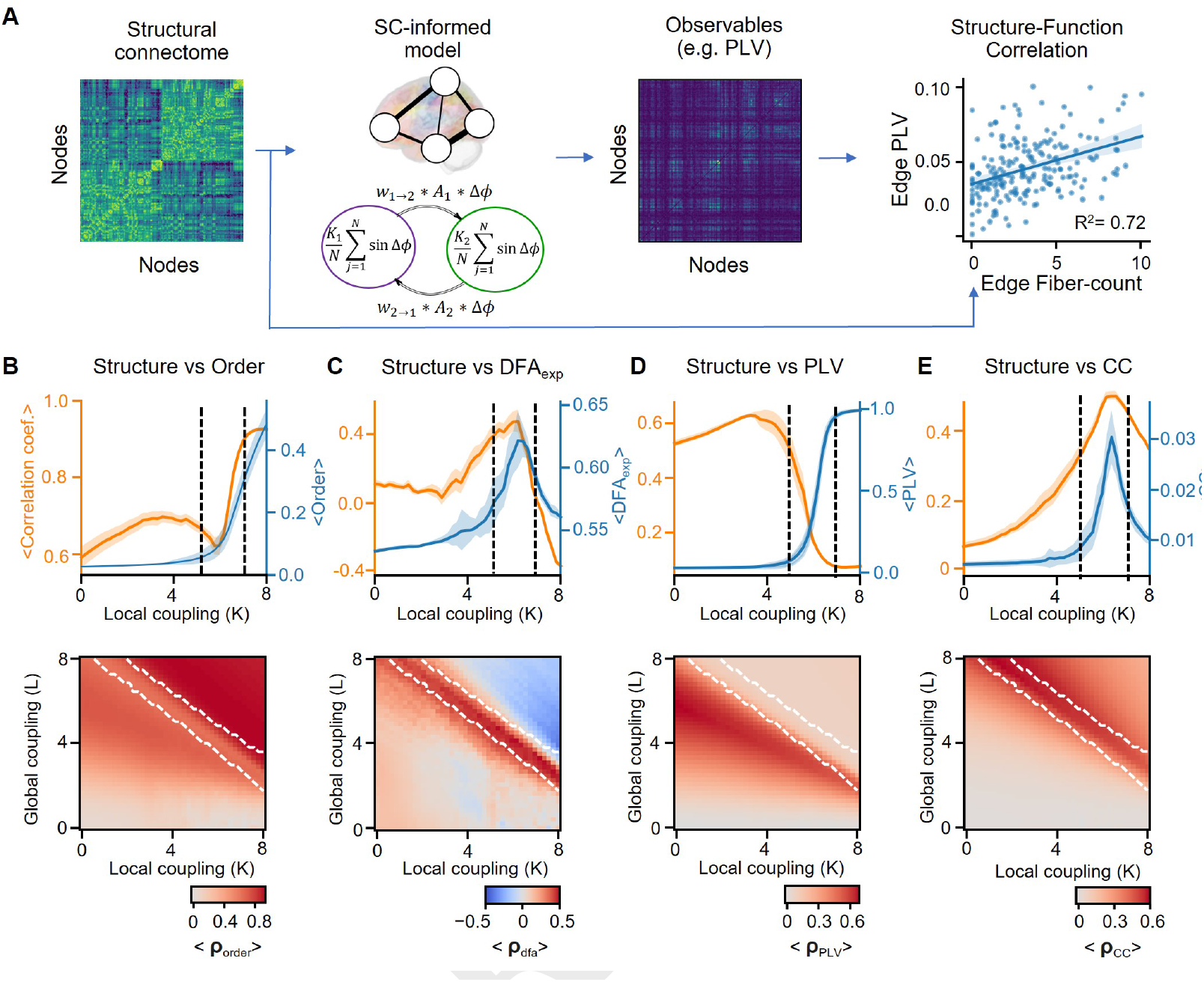
Modeling reveals diverse forms of Structure-Function coupling around the phase transition. **A** Analysis pipeline: The SC matrix of an individual is used to construct an SC-informed Kuramoto model, where edge coupling weights are derived from the SC. The model generates oscillatory dynamics, from which observables such as Phase Locking Value (PLV), DFA exponent (*DFA*_*exp*_), order, and amplitude cross-correlation (CC) are computed. Structure-function coupling is quantified as the correlation between these observables and structural measures (e.g., node strength or edge fiber count).**B-E** Correlation between model order and SC node strength (**B**), *DFA*_*exp*_ and SC node strength (**C**), PLV and SC edge weight (**D**), node order cross-correlation and edge weight (**E**) as a function of local coupling strength (K). Orange line indicates Pearson correlation coefficient (*ρ*) and the blue line average observable. Shaded areas represent confidence intervals (CIs) based on standard error (SE). On the bottom, heatmaps showing the Pearson correlation coefficient between the respective observables and structural measures (node strength or edge strength) as functions of local (K) and global (L) coupling strengths. The white contour outlines the critical regime consistent with Figure 2.

Investigating the structure-function correlations in the model at the individual node level, we found that node order was positively correlated with node strength (NS, average edge weight of each node), increasing slightly from the subcritical to the critical region, dipping at criticality, and then increasing sharply in the supercritical region (Figure 3B). Notably, this correlation remained close to zero independent of the local coupling in a region with low but non-zero global coupling parameter (Figure 3C).

The correlation between the DFA exponent and NS, on the other hand, fluctuated around zero in the subcritical region, peaked at criticality, and became strongly negative in the supercritical zone (Figure 3D). A similar phenomenon was observed on the KL surface where DFA vs. structure correlation was maximized in the critical region (Figure 3E). At the edge level, the correlation between edge strength and CC resembled the patterns observed with the DFA exponent, peaking at criticality while remaining low in both subcritical and supercritical zones (Figure 3H,I). The correlation between PLV and edge strength exhibited a unimodal shape, with a wide peak on the subcritical side and a drop around the critical peak (Figure 3F). Unlike other observables, PLV showed higher correlation with SC in regions where global coupling was stronger than local coupling (Figure 3G).

These results showed that the synchronization and criticality in oscillatory dynamics exhibited distinct and unique correlation regimes between structural and functional properties. In the critical region, the correlation between edge weights and intra-/inter-areal synchronization dipped but peaked for LRTCs and cross-correlation.

### Brain dynamics during the resting-state is the most correlated with observables in subcritical side of the extended critical regime

The model demonstrated a wide spectrum of critical dynamics across the transition from sub-to supercritical phases. We asked how similar the model dynamics was to the observed in human resting-state MEG recordings. Using MEG data, we computed key observables of oscillatory dynamics, including the weighted Phase Lag Index (wPLI), DFA exponents, and amplitude cross-correlation (Figure 4A). For wPLI and CC we also included in the analysis the average value across edges that belonged to the same node (node strength).

**Fig. 4.**
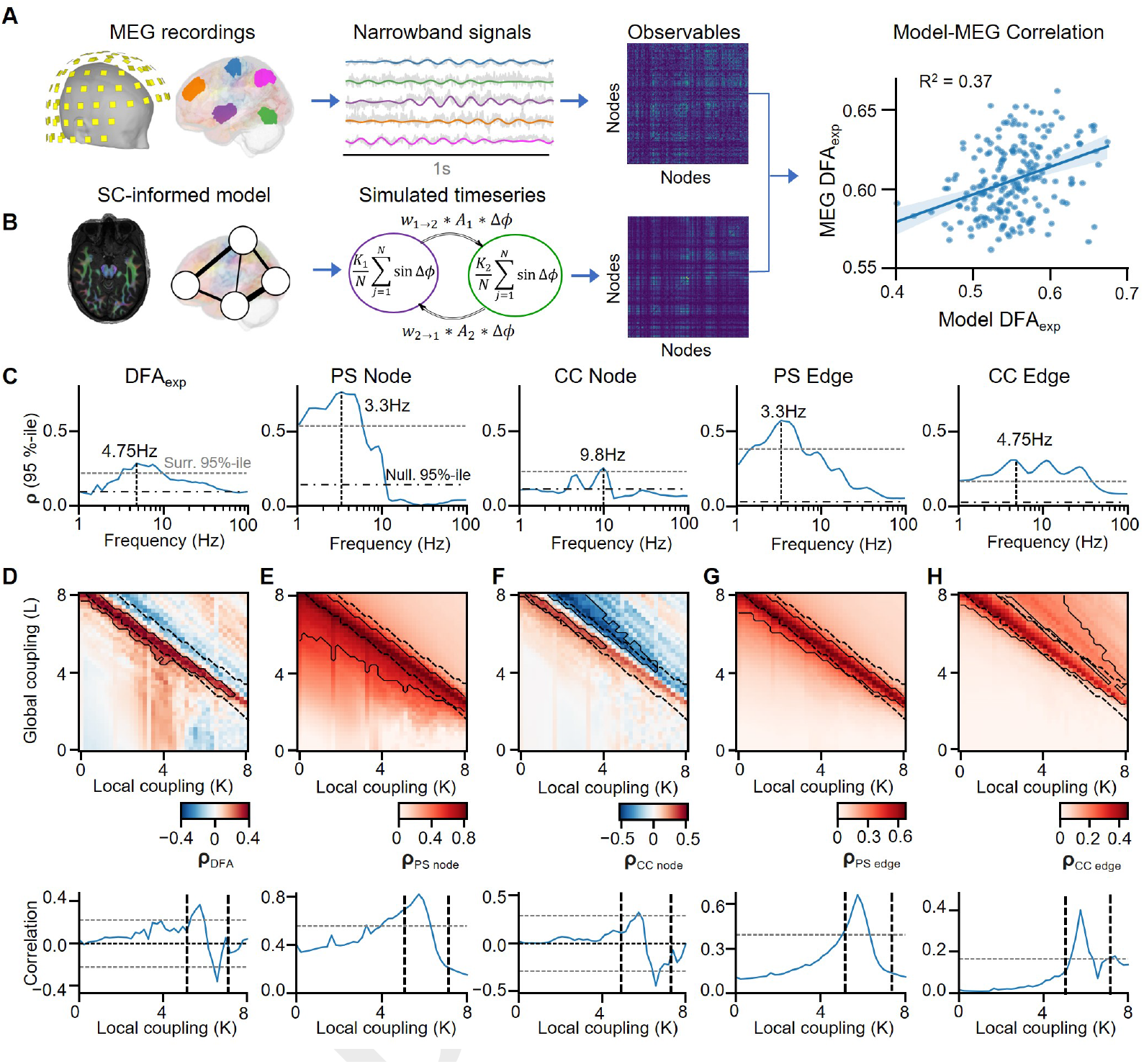
The models observables on the subcritical side of the extended critical regime most closely match the patterns seen in human MEG recordings. **A**. MEG analysis pipeline: MEG recordings are source-modeled to derive parcel-level time series, filtered into narrowband signals (e.g., using Morlet wavelets), and analyzed to compute observables such as phase synchronization matrices. **B** Modeling pipeline: The SC-informed Kuramoto model is used to simulate time series, from which the same observables (e.g., phase synchrony, DFA exponent) as in MEG data are computed. Pearson correlations are calculated between MEG and model observables at the parcel (node) level to quantify structure-function similarity. **C** The 95th percentile of the Pearson correlation coefficient across the KL-surface between MEG and model observables as a function of frequency. The dashed horizontal line indicates the statistical significance threshold determined via spin-permutation tests. The dash-dotted line indicates the 95th percentile of correlations in null model obtained by shuffling the structural connectome. The vertical dashed line highlights the frequency at which the correlation peaks. **D-H** Correlation maps and KL-surface slices for observables at the peak frequency identified in **C**. The Pearson correlation coefficient between simulated and MEG-derived DFA exponent (**E**), average phase-synchrony of a node (**D**), average cross-correlation of a node (**F**, edge phase synchrony (**G**) and edge cross-correlation (**H**) as a function of local (K) and global (L) coupling coefficients for frequency with maximum 95th percentile (see **C**). Black thick contours in the heatmaps indicate regions with statistically significant correlations (*p <* 0.01) based on spin-permutation tests. Black dashed contours delineate the critical regime (as defined in Figure 2). The line plots at the bottom represent a slice of the KL-surface for the same L = 4.

Next, we computed the same observables for simulated data but operationalized phase synchronization with PLV instead of wPLI (Figure 4B). Finally, to evaluate the similarity between real MEG recordings and simulated data, we estimated Pearson’s correlation coefficients between observables from model nodes/edges and MEG parcels/edges corresponding to the same anatomical regions.

We first assessed the similarity between model and MEG dynamics as a function of frequency. The model showed the most similar dynamics to the low-frequency activity, particularly in the theta frequency range (3–8 Hz) for the DFA exponent and node- and edge-level phase-synchrony (Figure 4C). However, the amplitude correlation exhibited several significant peaks including at theta (4.2 Hz), alpha (12.3 Hz), and beta frequencies (28.3 Hz).

To assess whether the similarities were a product of the realistic structural connectome, we computed the same model-MEG correlations for data simulated with two null models: using the shuffled structural connectome and a random uniform. For these null models, the correlations were nonsignificant (Figure 4C) showing that realistic architecture of node connectivity was crucial to produce in-vivo-like dynamics.

Second, we asked how the similarity between simulations and MEG behaved as a function of model control parameters. For each observable, we investigated the frequency with the highest correlation between the model and MEG data. We found that the significant correlation for all observables included in the analysis was found along the critical ridge (Figure 4D-H), which supports the notion that the human brain operates primarily on a subcritical side of the extended critical regime. Interestingly, the correlation coefficient for the DFA exponent and node-level CC was positive on the subcritical side of the critical regime but became negative on the critical-supercritical side (Figure 4D,F) which could indicate that the critical region in the human brain might be wider than observed in the model.

### The spectral properties of oscillatory activity are shaped by the distribution of underlying frequencies

Traditionally, oscillations are quantified by their magnitude, often using power spectral methods. Different system properties, such as interaction delays (52) or excitation-inhibition balance (69), have been shown to regulate the shape of the spectra and the emergence of oscillatory modes. However, the role of a system operating point remains an open question.

To replicate individual power spectral properties in the model, we aligned the oscillator frequency distribution with the shape of the recorded power spectrum. We achieved this by first computing the power spectrum density (PSD) of the MEG recordings removing the 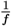 component using the FOOOF method (70). The resulting PSD was then transformed into a probability distribution of oscillator frequencies by normalizing it with the cumulative sum. For each node in the model, we sampled brain-region oscillator frequencies from these distributions individually for each MEG parcel (Figure 5A).

**Fig. 5.**
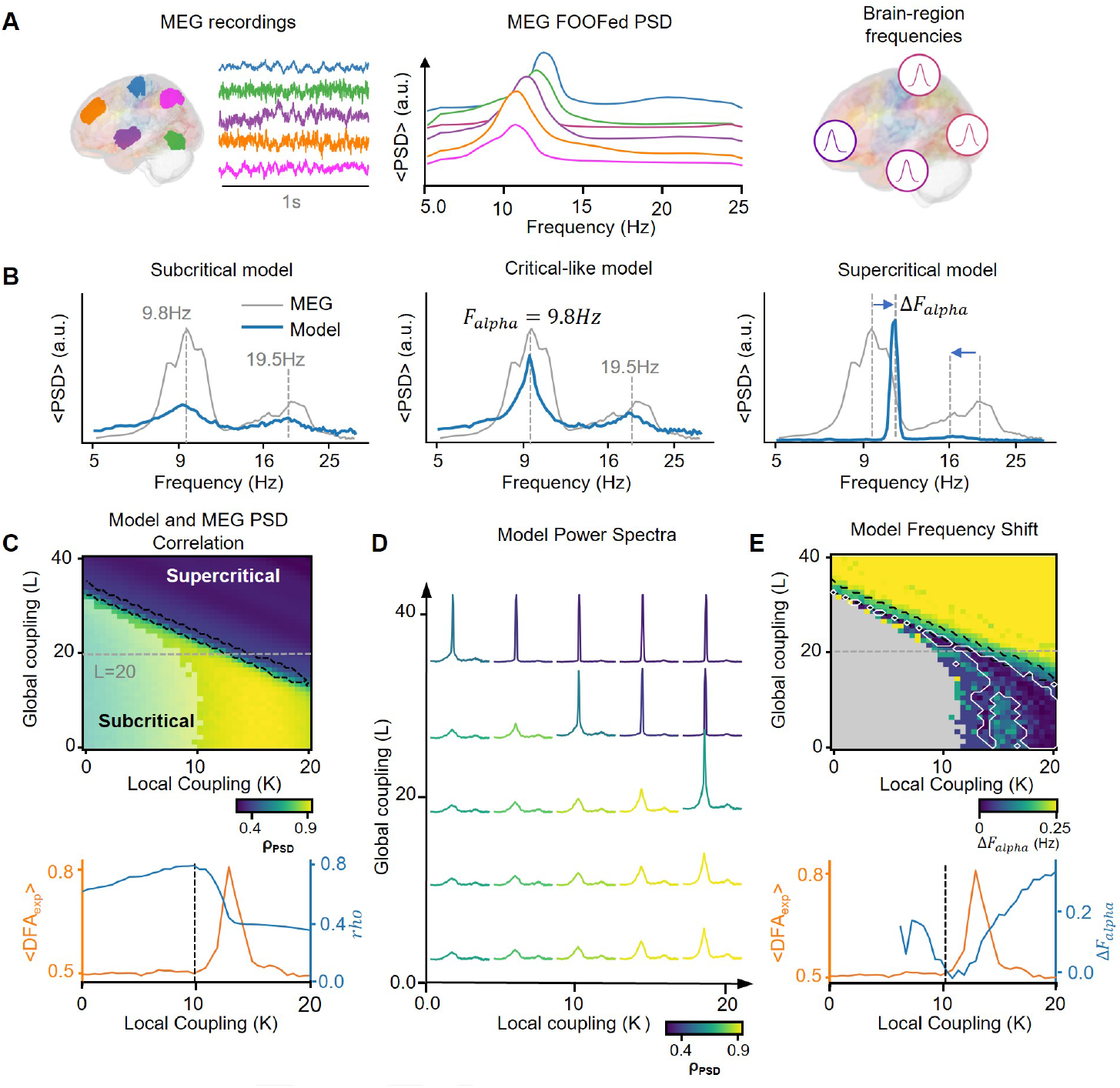
Critical-like state supports multi-frequency spectral properties. **A** Oscillator frequency distribution derived from parcel-level power spectral density (PSD) of real MEG recordings (left panel), with the 1*/f* component subtracted (middle panel) and are individually set for mode node that corresponds a brain area (right panel). **B** Average PSD across model nodes (blue) compared with MEG-derived PSD (gray) for three regimes: subcritical, critical, and supercritical. Dashed vertical lines indicate the alpha (*∼*9.8 Hz) and beta (*∼*19.5 Hz) frequency peaks. The critical model aligns closely with MEG-derived PSD, while subcritical and supercritical regimes deviate. **C** Pearson correlation coefficient between MEG-derived and simulated PSDs as a function of local (K) and global (L) coupling strengths (upper panel). The black contour highlights the critical ridge, defined by DFA exponent (*DFA*_*exp*_ *≥* 0.65). The semi-transparent area indicates regions without significant spectral peaks. Bottom panel: Correlation coefficient (orange) and average *DFA*_*exp*_ (blue) as functions of local coupling (K) for a fixed global coupling (L = 20). **D**Model PSDs for different combinations of local (K) and global (L) coupling strengths, color-coded by their correlation with MEG-derived PSDs, as shown in **C**. Simulated spectra with critical-like dynamics exhibit the most realistic multi-peak structure. **E**. Difference between simulated and MEG alpha peak frequencies (Δ*F*_*alpha*_) averaged across model nodes as a function of control parameters. The white contour highlights regions with significant alignment of alpha peaks. Bottom panel: Δ*F*_*alpha*_ (orange) and average *DFA*_*exp*_ (blue) as functions of local coupling (K), indicating the region of optimal alpha frequency alignment with critical-like dynamics.

To assess the similarity between the real and simulated data, we computed the PSD of the simulated and MEG data using the Welch method and estimated the Pearson correlation coefficient between them. In addition, we estimated the model frequency shift calculated by the absolute distance between the alpha peaks in the model and the MEG spectra.

Initially, we found that regions with low intra-nodal coupling and high inter-nodal coupling in the subcritical side showed low, non-significant peaks in the power spectrum. In the supercritical model, a dominant peak appeared in the power spectra at the alpha band. Additionally, the power spectrum exhibited a slight drift towards the weighted average of alpha (9.8 Hz) and beta (19.5 Hz) frequencies, calculated as 12.85 Hz weighted by frequency probability. In contrast, the spectrum of the critical-like model was well-matched, with the peaks corresponding to those observed in the real data (Figure 5B).

Since the properties of critical-like dynamics, particularly the critical state’s position within the control parameter space, depend on the oscillator frequency distribution, we initially investigated whether a model with a heterogeneous non-parametric frequency distribution would exhibit such dynamics. Our findings confirmed that a model initialized with a non-parametric distribution of oscillator frequencies did indeed display critical-like dynamics and LRTCs emerged during the phase transition at the same time preserving the multi-frequency activity (see Supplementary Figure 3).

Analyzing the similarity between the model and the MEG-derived power spectrum, we found that the highest correlation was achieved on the subcritical side of the extended critical regime, with high local and low global coupling strength, and decayed after the critical ridge (Figure 5C,D).

As the model transitioned from the subcritical to the critical regime, oscillatory peaks began to emerge. These peaks, largely absent at low control parameters, grew with increasing coupling strength and became dominated by the alpha band near the critical zone. Interestingly, the peaks were well-aligned with the real data in the subcritical and critical states, but in the supercritical regime, the alpha and beta frequency peaks exhibited a small drift toward the weighted average. The low correlation between the simulated and real data could be explained by this drift in the power spectrum properties (Figure 5E).

These findings suggest that operating near criticality achieved a precise balance in multi-modal spectral properties. This balance enabled the model to generate significant peaks in the power spectrum, closely reproduce MEG-derived PSDs, and prevent collapse into a singular frequency maintaining a bimodal spectrum.

## Discussion

We advance here a Hierarchical Kuramoto model that captures concurrently both functional-connectivity and brain-criticality phenomena and enables linking them across meso- and macroscopic scales of brain synchronization dynamics. Synchronization and criticality are interconnected in emergent brain activity so that their *in-vivo* inter-individual and - regional variability is likely driven by variability in the underlying individual and local operating points, respectively (21). Thus, a generative model that integrates these two aspects is essential for studying brain dynamics and developing experimentally testable, mechanistic predictions. Upon increasing local or inter-areal coupling, the model exhibited a transition from an asynchronous to a synchronous phase with emergent local and inter-areal critical-like dynamics at the phase transition across an extended critical regime (21, 63, 71).

We assessed the plausibility of the model with three lines of inquiries. First, we discovered that the structure-function coupling in the model was both strongly dependent on the operating point in an opposing manner for phase synchronization and amplitude dynamics, which suggests that different facets of functional connectivity exhibit distinct modes of structurefunction coupling in human brain networks. Next, assessing the anatomical similarity of the model observables with those observed with resting-state MEG revealed moderate to large correlations that peaked in the subcritical-side of the model’s extended critical regime across all observables. These correlations vanished in the supercritical side of the critical regime as well in the sub- and supercritical phases, suggesting that human brains predominantly operate on the subcritical side of an extended critical regime rather than at peak criticality. Finally, the hierarchical organization of the model enabled it to exhibit local oscillations concurrently at multiple frequencies, as found *in-vivo*, and addressing their dependence on the operating point *in-silico*. Modeling showed that multi-frequency oscillations peaked at criticality and were most similar with MEG observations again in the subcritical side of the extended critical regime. These findings establish a new modeling framework for assessing how local and global operating points regulate emergent brain dynamics, opening new avenues for mechanistic understanding of macroscale brain activity.

### Concurrent modeling of local and global critical synchronization dynamics

Brain dynamics arise through scales from microcircuits to macroscopic populations and thus concurrent consideration of local and inter-areal dynamics is essential for modeling brain activity. Hierarchical methods have recently gained traction in modeling of neuronal activity, where they have been used to combine models at different scales (72) such as both spiking and mean-field activity simultaneously to accurately predict an effect of deep brain stimulation (73). It has also been used to simulate local and networklevel interactions in the Digital Brain Twin models (74) or multiple populations, in particular, two nodes of spiking neuron populations have been used to assess the effects of local network dynamics on inter-node amplitude- and phase-coupling (60) and a four-node-model of Kuramoto oscillators has been used to model inter-node cross-frequency synchronization dynamics (50). Segregated populations of ‘excitatory’ and ‘inhibitory’ oscillators have also been developed to evaluate the role of the excitation–inhibition balance in Kuramoto synchronization dynamics (67).

Extending a hierarchical implementation of the Kuramoto model to the whole-brain scale, we found here how simultaneous modeling of local and inter-areal synchronization dynamics reveals a multi-scale transition from an asynchronous to a synchronized phase in both node order and inter-areal phase synchrony. At the node level, the phase transition was associated with emergence of long-range temporal correlations (LRTCs) that evidence local critical-like dynamics (41, 75, 76). In inter-areal coupling, this phase transition in synchronization was associated with maximized node-order correlations, i.e., amplitude correlations of the local oscillations. While in the model node order/amplitude is directly determined by local synchronization, in fact, this is also the case in MEG and EEG where the local signal magnitude is primarily determined by local synchronization (77, 78). Amplitude correlations in experimental data that largely reflect inter-areal coupling of local synchronization fluctuations. Together with phase synchrony (3, 5, 11, 21), amplitude correlations are a key mode of “functional connectivity” in human brain dynamics both at macro- (79) and meso-scopic levels (2). While the structures phase-synchrony and amplitude-correlation networks have considerable anatomical similarity (80–82), the model predicts that they are mechanistically distinct so that phase synchrony is monotonically dependent on the operating point while amplitude correlations exhibit a quadratic dependence. These findings extend to the wholebrain level observations made with a two-node system of spiking neurons (60) as well as with maximization of dynamic correlations at criticality in physical models, such as the Ising model (32).

### Unique profiles of structure-function coupling across the critical phase transition

Large-scale brain activity is shaped and constrained by inter-areal structural connectivity (SC) defined especially by white-matter axonal projections (83, 84). Both fMRI and MEG studies have shown that functional connectivity networks are correlated with the SC networks but with notable anatomical variability in this structurefunction coupling (SFC) (85, 86) so that SFC is stronger in sensorimotor cortices than in association areas. The relationship of SFC with brain criticality and the operating point of neuronal systems has, however, remained controversial. On one hand, computational modeling suggests that SC and FC decouple at criticality due to the emergence of correlations (functional connectivity) between areas that are not directly structurally connected (39). On the other hand, comparisons between modeled and fMRI resting state networks, suggest that FC emerges directly from SC at criticality (68, 87).

To address this conundrum, we assessed the anatomical similarity between SC and oscillatory dynamics as a function of the model’s operating point. Synchronization was strongly correlated with SC in the subcritical regime, but this relationship broke down at criticality as a result of emergent indirect correlations. Thus while SC constrains phase synchronization in the subcritical phase, even nodes with weak SC can synchronize via higher-order interactions at criticality.

However, in contrast with phase synchronization, both the local DFA exponents and inter-areal amplitude correlations exhibited the strongest correlations with SC at criticality. The model thus makes opposing and experimentally predictions for the SFC of phase and amplitude coupling along the critical regime (21).

SFC has important implications for the controllability of the brain as complex system. The differences in the network structure support distinct roles in the control of brain states, with strongly connected hub regions typically serving as the most impactful control points (88, 89). However, rather than operating to a single regime of dynamics, neuronal systems exhibit multiple distinct dynamical states (90, 91). Linear controllability has been assessed in a series of studies and approaches to brain modeling (88, 92, 93). The present _1_ findings, showing highly non-linear dynamics and emergent second-order interactions at criticality, suggest that in the critical regime linear control approaches might be less effective than in the subcritical state (21).

### Structures of model functional networks match MEG data in the subcritical side of an extended critical regime

The “critical brain” hypothesis suggests that the brain operates at intermediate levels of synchronization near a phase transition between order and disorder (24, 25) where power-law temporal correlations (21, 41, 75) and neuronal avalanches (41, 94, 95) emerge. However, the operating point of human brains in vivo is has remained an increasingly central unresolved objective in the field (21, 96). Comparing computational model and experimental data is a well-established way to understand the causal effects of structure and mechanisms on emergent dynamics in complex systems (46). Human whole-brain scale modeling has shown that models tuned to criticality (40, 68, 97), those operating at maximum metastability (52, 98, 99), as well as the critical-Hopf-bifurcation model (58) achieve the greatest similarity with MEG- and fMRI-based experimentally observed functional connectivity. In addition, the distribution of DFA exponents observed in CROS model at criticality was similar to human MEG data observations (61). On the other hand, earlier studies have not assessed the model-experiment correlations simultaneously for FC and criticality while both are dependent on the operating point. To address this, we compared the topographies of model observables with those in resting-state MEG. Throughout all studied observables, the greatest similarity between model and experimental observables was found systematically on the subcritical side of an extended critical regime. These findings extend the prior modeling and experimental studies to yield a new line of evidence for the notion that most human individuals during resting-state operate on the subcritical side rather than at the peak or in the supercritical side of the critical regime (21).

### Criticality balances multi-frequency activities in a system

The human brain exhibits oscillations at multiple frequencies concurrently (6, 50, 100, 101) which are typically operationalized as peaks in the power spectrum (70). Prior studies have indicated that specific aspects of the power spectrum are influenced by factors such as the E/I balance (69, 102), delayed interactions (52), or cortical layers’ organization (103). However, typically models with only a single central frequency has been studies and the role of operating point in a system with several concurrent oscillation frequencies has remained unknown.

When we initialized the model to exhibit concurrent oscillations at multiple frequencies, similarly to in-vivo observations, multiple peaks in the power spectrum emerged in the subcritical-to-critical state, most closely resembling MEG-derived data. In the supercritical phase, we observed a single oscillatory mode becoming dominant with its frequency shifted towards the weighted average of individual peak frequencies, whereas for in subcritical regime with low local coupling no significant spectral peaks were observed.

Our results show that the spectral properties are defined by both the distribution of underlying oscillators’ frequencies and the model’s critical dynamics. The shape of the PSD was primarily dictated by the frequency distribution of the oscillators. When this distribution aligned with the shape of the MEG-derived PSD, the model was able to reproduce key properties such as location of spectral peaks. The operating point regulates the emergence of multiple peaks, from an almost complete absence of oscillations in the subcritical region to a singular frequency in the supercritical regime. Notably, in the subcritical-to-critical regime, the model preserves a multi-modal spectrum and achieves the highest similarity with PSD as observed in MEG experiments. Based on these results, we propose that operating in this regime provides sufficient local synchronization to produce sustained oscillations, as indicated by significant peaks in the power spectrum, while also preserving the necessary variability in activity to support multi-frequency dynamics.

## Conclusion

Criticality is a rapidly emerging framework in brain research that posits an operating point governing the variability in neuronal dynamics. However, the causal effects of this operating point on functional properties has remained unclear. In this work, we introduced a generative hierarchical framework for modeling critical-like dynamics that explicitly captures both local and inter-areal synchronization. Our model exhibits emergent multi-scale dynamics that replicate several features of critical meso- and macro-scale brain activity observed in electrophysiological recordings. The results demonstrated the causal effect of the operating point on structure–function coupling and spectral properties of multi-frequency oscillations in the model. We believe this new approach highlights the importance of critical-like state in the complex dynamics and opens new avenues for wholebrain modeling to investigate the causal impact of control parameters on brain activity to guide personalized treatment strategies, such as targeted brain stimulation.

## Materials and Methods

### Hierachical kuramoto model

The classical Kuramoto model describes the synchronization dynamics of coupled oscillators, where each oscillator aligns its phase with others based on coupling strength

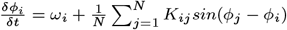

where *N* is the total number of oscillators, *ϕ*_*i*_ is the i-th oscillator phase, *ω*_*i*_ is the frequency of the i-th oscillator, and *K*_*ij*_ is the coupling parameter between the i-th and j-th oscillators.

In this framework, each oscillator represents a brain region or parcel, and the coupling parameter reflects the strength of SC between two zones. However, a single Kuramoto oscillator has constant amplitude in complex domain, which makes comparison with real brain imaging data a challenging task. To overcome this, we introduce a hierarchical extension of the Kuramoto model with multiple nodes comprising multiple locally coupled oscillators. This corresponds to the analysis of electrophysiological recordings, where each electrode or cortical parcel represents a large number of locally interacting neurons. Our model thus allows for the representation of both local synchronization within nodes and long-range inter-areal interactions between nodes. The behavior of individual oscillators within a node is governed by three terms:

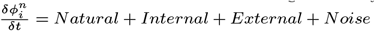

Where 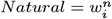 represents the central frequency of the i-th oscillator in the n-th node and 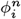 is the phase of the i-th oscillator in the n-th node,

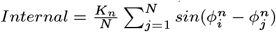

where *Kn* is the local coupling parameter of the n-th node and N is the total number of oscillators within the node. In this context, the phase of each oscillator is shifted towards the average phase of oscillators within a node,

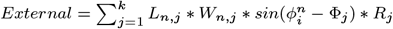

Where *L*_*n,j*_ is the coupling coefficient between the n-th and j-th nodes (global control parameter), *W*_*n,j*_ is the SC between the n-th and j-th nodes, Φ_*j*_ is the cyclic average of the phases of the n-th node defined as 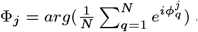 and *R* is the order of the j-th node (see the definition below). In this context, the phase of each oscillator is compared to the average phase of other nodes and weighted by a node order and edge strength.

and 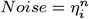 is the white noise for the i-th oscillator in the n-th node.

Although trigonometric functions are costly to compute and we used the complex representation to improve computational performance, we used a complex-valued representation for oscillator states. Thus, the coupling function between oscillators x and y becomes *imag*(*x* y*^*\**^), where x and y are complex values representing an oscillator phase, and *y*^*\**^ is a conjugate of y.

## Computational experiments

We simulated 6 minutes of restingstate activity for each combination of control parameters. In the experiments for Figures 2, 3, and 4, the parameters K and L took on equally spaced values from 0 to 8. In the experiment for Figure 5, the local coupling was varied from 0 to 20, and the global coupling from 0 to 40. We removed the first minute of data of each simulation which can be seen as a “warm-up period” to avoid inflated inflated DFA exponent values. The detailed explanation and visualization of model parameters and their values for each experimenmt are shown in Supplementary Figure 4 and Supplementary Table 1.

### Observables of oscillatory dynamics

We quantified local synchrony with the order parameter, defined as the absolute value of the average complex phase across oscillators in each node. Inter-areal phase interactions were estimated using the phase locking value (PLV) for the model and the weighted Phase Lag Index (wPLI, (104, 105)) for MEG data. Long-range temporal correlations (LRTCs) were measured using Detrended Fluctuation Analysis (DFA, (76, 106)) computed in the Fourier domain with robust linear regression. Detailed methodology is provided in the Supplementary Information.

### Data, materials and software availability

The model and experimental code will be open-sourced following publication.

### Data acquisition

The study protocol for MEG, MRI, and DWI data was approved by the HUS ethical committee (**HUS/3043/2021**, 27.4.2022), written informed consent was obtained from each participant prior to the experiment, and all research was carried out according to the Declaration of Helsinki. The details of MEG and DWI data recording processing can be found in the Supplementary Information.

## ACKNOWLEDGMENTS

We thank Sheng Wang, Felix Siebenhühner, and Joonas J. Juvonen for feedback about the study and their input on the manuscript. We also thank professor Guido Nolte on helpful comments on an earlier version of the manuscript. The simulations have been done on the LUMI supercomputer. This study was supported by the Academy of Finland (J.M.P., project numbers: 296304), by the Juselius Foundation (S.P and J.M.P project number 240156).

## Notes

### Competing Interest Statement

The authors have declared no competing interest.

### Summary of Updates

We revised the text and the figures

## References

1. G Buzsáki, CA Anastassiou, C Koch, The origin of extracellular fields and currents–EEG, ECoG, LFP and spikes. Nat. Rev. Neurosci. 13, 407–420 (2012).

2. G Arnulfo, J Hirvonen, L Nobili, S Palva, JM Palva, Phase and amplitude correlations in resting-state activity in human stereotactical eeg recordings. NeuroImage 112, 114–127 (2015).

3. G Arnulfo, et al., Long-range phase synchronization of high-frequency oscillations in human cortex. Nat. Commun. 11 (2020).

4. N Williams, et al., The influence of inter-regional delays in generating large-scale brain networks of phase synchronization. NeuroImage 279, 120318 (2023).

5. S Palva, JM Palva, Discovering oscillatory interaction networks with m/eeg: challenges and breakthroughs. Trends Cogn. Sci. 16, 219–230 (2012).

6. JM Palva, S Palva, K Kaila, Phase synchrony among neuronal oscillations in the human cortex. J. Neurosci. 25, 3962–3972 (2005).

7. E Glerean, J Salmi, JM Lahnakoski, I. Jääskeläinen, M Sams, Functional magnetic resonance imaging phase synchronization as a measure of dynamic functional connectivity. Brain Connect. 2, 91–101 (2012).

8. P Fries, Rhythms for cognition: Communication through coherence. Neuron 88, 220–235 (2015).

9. M Vinck, et al., Principles of large-scale neural interactions. Neuron 111, 987–1002 (2023).

10. M Schneider, et al., A mechanism for inter-areal coherence through communication based on connectivity and oscillatory power. Neuron 109, 4050–4067.e12 (2021).

11. JM Palva, S Monto, S Kulashekhar, S Palva, Neuronal synchrony reveals working memory networks and predicts individual memory capacity. Proc. Natl. Acad. Sci. 107, 7580–7585 (2010).

12. H Haque, M Lobier, JM Palva, S Palva, Neuronal correlates of full and partial visual conscious perception. Conscious. Cogn. 78, 102863 (2020).

13. F Siebenhühner, SH Wang, JM Palva, S Palva, Cross-frequency synchronization connects networks of fast and slow oscillations during visual working memory maintenance. eLife 5 (2016).

14. SH Wang, et al., Neuronal synchrony and critical bistability: Mechanistic biomarkers for localizing the epileptogenic network. preprint (2023).

15. LI Boon, et al., A systematic review of meg-based studies in parkinson’s disease: The motor system and beyond. Hum. Brain Mapp. 40, 2827–2848 (2019).

16. S Pusil, et al., Hypersynchronization in mild cognitive impairment: the ‘x’model. Brain 142, 3936–3950 (2019).

17. G Alamian, et al., Alterations of intrinsic brain connectivity patterns in depression and bipolar disorders: A critical assessment of magnetoencephalography-based evidence. Front. Psychiatry 8 (2017).

18. S Zhang, et al., Association between abnormal default mode network activity and suicidality in depressed adolescents. BMC Psychiatry 16 (2016).

19. RH Kaiser, JR Andrews-Hanna, TD Wager, D. Pizzagalli, Large-scale network dysfunction in major depressive disorder: A meta-analysis of resting-state functional connectivity. JAMA Psychiatry 72, 603 (2015).

20. Y Mohammadi, MH Moradi, Prediction of depression severity scores based on functional connectivity and complexity of the eeg signal. Clin. EEG Neurosci. 52, 52–60 (2020).

21. M Fuscà, et al., Brain criticality predicts individual levels of inter-areal synchronization in human electrophysiological data. Nat. Commun. 14 (2023).

22. S Palva, S Monto, JM Palva, Graph properties of synchronized cortical networks during visual working memory maintenance. NeuroImage 49, 3257–3268 (2010).

23. C Micou, T O’Leary, Representational drift as a window into neural and behavioural plasticity. Curr. Opin. Neurobiol. 81, 102746 (2023).

24. WL Shew, D Plenz, The functional benefits of criticality in the cortex. The Neurosci. 19, 88–100 (2012).

25. DR Chialvo, Emergent complex neural dynamics. Nat. Phys. 6, 744–750 (2010).

26. K Linkenkaer-Hansen, VV Nikouline, JM Palva, RJ Ilmoniemi, Long-range temporal correlations and scaling behavior in human brain oscillations. The J. Neurosci. 21, 1370–1377 (2001).

27. JM Beggs, D Plenz, Neuronal avalanches in neocortical circuits. The J. Neurosci. 23, 11167–11177 (2003).

28. N Lotfi, et al., Statistical complexity is maximized close to criticality in cortical dynamics. Phys. Rev. E 103 (2021).

29. O Kinouchi, M Copelli, Optimal dynamical range of excitable networks at criticality. Nat. Phys. 2, 348–351 (2006).

30. V Zimmern, Why brain criticality is clinically relevant: A scoping review. Front. Neural Circuits 14 (2020).

31. K Heiney, et al., Criticality, connectivity, and neural disorder: A multifaceted approach to neural computation. Front. Comput. Neurosci. 15 (2021).

32. JM Beggs, The criticality hypothesis: how local cortical networks might optimize information processing. Philos. Transactions Royal Soc. A: Math. Phys. Eng. Sci. 366, 329–343 (2007).

33. SS Poil, R Hardstone, HD Mansvelder, K Linkenkaer-Hansen, Critical-state dynamics of avalanches and oscillations jointly emerge from balanced excitation/inhibition in neuronal networks. J. Neurosci. 32, 9817–9823 (2012).

34. WL Shew, H Yang, T Petermann, R Roy, D Plenz, Neuronal avalanches imply maximum dynamic range in cortical networks at criticality. The J. Neurosci. 29, 15595–15600 (2009).

35. J Simola, et al., Genetic polymorphisms in comt and bdnf influence synchronization dynamics of human neuronal oscillations. iScience 25, 104985 (2022).

36. JM Shine, et al., Computational models link cellular mechanisms of neuromodulation to large-scale neural dynamics. Nat. Neurosci. 24, 765–776 (2021).

37. JM Shine, Neuromodulatory control of complex adaptive dynamics in the brain. Interface Focus. 13 (2023).

38. RE Brown, JT McKenna, Turning a negative into a positive: Ascending gabaergic control of cortical activation and arousal. Front. Neurol. 6 (2015).

39. JM Beggs, N Timme, Being critical of criticality in the brain. Front. physiology 3, 163 (2012).

40. M Rubinov, O Sporns, JP Thivierge, M Breakspear, Neurobiologically realistic determinants of self-organized criticality in networks of spiking neurons. PLoS Comput. Biol. 7, e1002038 (2011).

41. JM Palva, et al., Neuronal long-range temporal correlations and avalanche dynamics are correlated with behavioral scaling laws. Proc. Natl. Acad. Sci. 110, 3585–3590 (2013).

42. H Bruining, et al., Measurement of excitation-inhibition ratio in autism spectrum disorder using critical brain dynamics. Sci. Reports 10 (2020).

43. P Ritter, M Schirner, AR McIntosh, VK Jirsa, The virtual brain integrates computational modeling and multimodal neuroimaging. Brain Connect. 3, 121–145 (2013).

44. A Ponce-Alvarez, G Deco, The hopf whole-brain model and its linear approximation. Sci. Reports 14 (2024).

45. M Breakspear, S Heitmann, A Daffertshofer, Generative models of cortical oscillations: neurobiological implications of the kuramoto model. Front. human neuroscience 4, 190 (2010).

46. M Breakspear, Dynamic models of large-scale brain activity. Nat. Neurosci. 20, 340–352 (2017).

47. G Deco, et al., Resting-state functional connectivity emerges from structurally and dynamically shaped slow linear fluctuations. J. Neurosci. 33, 11239–11252 (2013).

48. A Daffertshofer, R Ton, B Pietras, ML Kringelbach, G Deco, Scale-freeness or partial synchronization in neural mass phase oscillator networks: Pick one of two? NeuroImage 180, 428–441 (2018).

49. ML Kringelbach, AR McIntosh, P Ritter, VK Jirsa, G Deco, The rediscovery of slowness: Exploring the timing of cognition. Trends Cogn. Sci. 19, 616–628 (2015).

50. F Siebenhühner, et al., Genuine cross-frequency coupling networks in human resting-state electrophysiological recordings. PLOS Biol. 18, e3000685 (2020).

51. P Tewarie, et al., Tracking dynamic brain networks using high temporal resolution meg measures of functional connectivity. NeuroImage 200, 38–50 (2019).

52. J Cabral, et al., Metastable oscillatory modes emerge from synchronization in the brain spacetime connectome. Commun. Phys. 5 (2022).

53. G Deco, et al., Awakening: Predicting external stimulation to force transitions between different brain states. Proc. Natl. Acad. Sci. 116, 18088–18097 (2019).

54. J Cabral, E Hugues, O Sporns, G Deco, Role of local network oscillations in resting-state functional connectivity. NeuroImage 57, 130–139 (2011).

55. JP Hobbs, JL Smith, JM Beggs, Aberrant neuronal avalanches in cortical tissue removed from juvenile epilepsy patients. J. Clin. Neurophysiol. 27, 380–386 (2010).

56. Y Liu, KA Dahmen, Unexpected universality in static and dynamic avalanches. Phys. Rev. E 79 (2009).

57. MJ Aburn, CA Holmes, JA Roberts, TW Boonstra, M Breakspear, Critical fluctuations in cortical models near instability. Front. Physiol. 3 (2012).

58. G Deco, ML Kringelbach, VK Jirsa, P Ritter, The dynamics of resting fluctuations in the brain: metastability and its dynamical cortical core. Sci. Reports 7 (2017).

59. JA Roberts, et al., Metastable brain waves. Nat. Commun. 10 (2019).

60. AE Avramiea, A Masood, HD Mansvelder, K Linkenkaer-Hansen, Long-range amplitude coupling is optimized for brain networks that function at criticality. J. Neurosci. 42, 2221–2233 (2022).

61. L Dalla Porta, M Copelli, Modeling neuronal avalanches and long-range temporal correlations at the emergence of collective oscillations: Continuously varying exponents mimic m/eeg results. PLOS Comput. Biol. 15, e1006924 (2019).

62. Y Kuramoto, International symposium on mathematical problems in theoretical physics. Lect. notes Phys. 30, 420 (1975).

63. P Villegas, P Moretti, MA Muñoz, Frustrated hierarchical synchronization and emergent complexity in the human connectome network. Sci. Reports 4 (2014).

64. DP Koller, M Schirner, P Ritter, Human connectome topology directs cortical traveling waves and shapes frequency gradients. Nat. Commun. 15 (2024).

65. A Escrichs, et al., Whole-brain turbulent dynamics predict responsiveness to pharmacological treatment in major depressive disorder. Mol. Psychiatry (2024).

66. SH Wang, et al., Critical-like brain dynamics in a continuum from second-to first-order phase transition. The J. Neurosci. 43, 7642–7656 (2023).

67. S Kuroki, K Mizuseki, Excitation–inhibition balance controls synchronization in a simple model of coupled phase oscillators. preprint (2024).

68. A Haimovici, E Tagliazucchi, P Balenzuela, DR Chialvo, Brain organization into resting state networks emerges at criticality on a model of the human connectome. Phys. Rev. Lett. 110, 178101 (2013).

69. F Lombardi, HJ Herrmann, L de Arcangelis, Balance of excitation and inhibition determines 1/f power spectrum in neuronal networks. Chaos: An Interdiscip. J. Nonlinear Sci. 27 (2017).

70. T Donoghue, et al., Parameterizing neural power spectra into periodic and aperiodic components. Nat. Neurosci. 23, 1655–1665 (2020).

71. MA Muñoz, Colloquium: Criticality and dynamical scaling in living systems. Rev. Mod. Phys. 90 (2018).

72. E D’Angelo, V Jirsa, The quest for multiscale brain modeling. Trends Neurosci. 45, 777–790 (2022).

73. JM Meier, et al., Virtual deep brain stimulation: Multiscale co-simulation of a spiking basal ganglia model and a whole-brain mean-field model with the virtual brain. Exp. Neurol. 354, 114111 (2022).

74. HE Wang, et al., Virtual brain twins: from basic neuroscience to clinical use. Natl. Sci. Rev. 11 (2024).

75. K Linkenkaer-Hansen, VV Nikouline, JM Palva, RJ Ilmoniemi, Long-range temporal correlations and scaling behavior in human brain oscillations. J. Neurosci. 21, 1370–1377 (2001).

76. R Hardstone, et al., Detrended fluctuation analysis: a scale-free view on neuronal oscillations. Front. physiology 3, 450 (2012).

77. VN Murthy, EE Fetz, Synchronization of neurons during local field potential oscillations in sensorimotor cortex of awake monkeys. J. Neurophysiol. 76, 3968–3982 (1996).

78. O Herreras, Local field potentials: Myths and misunderstandings. Front. Neural Circuits 10 (2016).

79. MJ Brookes, et al., Investigating the electrophysiological basis of resting state networks using magnetoencephalography. Proc. Natl. Acad. Sci. 108, 16783–16788 (2011).

80. A Zhigalov, G Arnulfo, L Nobili, S Palva, JM Palva, Modular co-organization of functional connectivity and scale-free dynamics in the human brain. Netw. Neurosci. 1, 143–165 (2017).

81. M Siems, M Siegel, Dissociated neuronal phase- and amplitude-coupling patterns in the human brain. NeuroImage 209, 116538 (2020).

82. F Siebenhühner, JM Palva, S Palva, Node centrality in meg resting-state networks co-varies with neurotransmitter receptor and transporter density. bioRxiv (2024).

83. JS Damoiseaux, MD Greicius, Greater than the sum of its parts: a review of studies combining structural connectivity and resting-state functional connectivity. Brain Struct. Funct. 213, 525–533 (2009).

84. P Hagmann, et al., White matter maturation reshapes structural connectivity in the late developing human brain. Proc. Natl. Acad. Sci. 107, 19067–19072 (2010).

85. L. Suárez, RD Markello, RF Betzel, B Misic, Linking structure and function in macroscale brain networks. Trends Cogn. Sci. 24, 302–315 (2020).

86. ZQ Liu, G Shafiei, S Baillet, B Misic, Spatially heterogeneous structure-function coupling in haemodynamic and electromagnetic brain networks. NeuroImage 278, 120276 (2023).

87. H Lee, et al., Relationship of critical dynamics, functional connectivity, and states of consciousness in large-scale human brain networks. NeuroImage 188, 228–238 (2019).

88. S Gu, et al., Controllability of structural brain networks. Nat. Commun. 6 (2015).

89. CW Lynn, DS Bassett, The physics of brain network structure, function and control. Nat. Rev. Phys. 1, 318–332 (2019).

90. C Tu, et al., Warnings and caveats in brain controllability. NeuroImage 176, 83–91 (2018).

91. G Deco, VK Jirsa, PA Robinson, M Breakspear, K Friston, The dynamic brain: From spiking neurons to neural masses and cortical fields. PLoS Comput. Biol. 4, e1000092 (2008).

92. P Srivastava, et al., Models of communication and control for brain networks: distinctions, convergence, and future outlook. Netw. Neurosci. 4, 1122–1159 (2020).

93. E Nozari, et al., Macroscopic resting-state brain dynamics are best described by linear models. Nat. Biomed. Eng. 8, 68–84 (2023).

94. A Zhigalov, G Arnulfo, L Nobili, S Palva, JM Palva, Relationship of fast- and slow-timescale neuronal dynamics in human meg and seeg. The J. Neurosci. 35, 5385–5396 (2015).

95. O Shriki, et al., Neuronal avalanches in the resting meg of the human brain. The J. Neurosci. 33, 7079–7090 (2013).

96. KB Hengen, WL Shew, Is criticality a unified set-point of brain function? bioRxiv (2024).

97. RP Rocha, et al., Recovery of neural dynamics criticality in personalized whole-brain models of stroke. Nat. Commun. 13 (2022).

98. G Deco, et al., Rare long-range cortical connections enhance human information processing. Curr. Biol. 31, 4436–4448.e5 (2021).

99. F Castaldo, et al., Multi-modal and multi-model interrogation of large-scale functional brain networks. NeuroImage 277, 120236 (2023).

100. A Capilla, et al., The natural frequencies of the resting human brain: An meg-based atlas. NeuroImage 258, 119373 (2022).

101. V Myrov, et al., Rhythmicity of neuronal oscillations delineates their cortical and spectral architecture. Commun. Biol. 7 (2024).

102. R Gao, EJ Peterson, B Voytek, Inferring synaptic excitation/inhibition balance from field potentials. NeuroImage 158, 70–78 (2017).

103. R Sanchez-Todo, et al., A physical neural mass model framework for the analysis of oscillatory generators from laminar electrophysiological recordings. NeuroImage 270, 119938 (2023).

104. SH Wang, et al., Hyperedge bundling: A practical solution to spurious interactions in MEG/EEG source connectivity analyses. NeuroImage 173, 610–622 (2018).

105. M Vinck, R Oostenveld, M van Wingerden, F Battaglia, CM Pennartz, An improved index of phase-synchronization for electrophysiological data in the presence of volume-conduction, noise and sample-size bias. NeuroImage 55, 1548–1565 (2011).

106. G Nolte, M Aburidi, AK Engel, Robust calculation of slopes in detrended fluctuation analysis and its application to envelopes of human alpha rhythms. Sci. Reports 9 (2019).

